# b-move: faster bidirectional character extensions in a run-length compressed index

**DOI:** 10.1101/2024.05.30.596587

**Authors:** Lore Depuydt, Luca Renders, Simon Van de Vyver, Lennart Veys, Travis Gagie, Jan Fostier

## Abstract

Due to the increasing availability of high-quality genome sequences, pan-genomes are gradually replacing single consensus reference genomes in many bioinformatics pipelines to better capture genetic diversity. Traditional bioinformatics tools using the FM-index face memory limitations with such large genome collections. Recent advancements in run-length compressed indices like Gagie et al.’s r-index and Nishimoto and Tabei’s move structure, alleviate memory constraints but focus primarily on backward search for MEM-finding. Arakawa et al.’s br-index initiates complete approximate pattern matching using bidirectional search in run-length compressed space, but with significant computational overhead due to complex memory access patterns. We introduce b-move, a novel bidirectional extension of the move structure, enabling fast, cache-efficient bidirectional character extensions in run-length compressed space. It achieves bidirectional character extensions up to 8 times faster than the br-index, closing the performance gap with FM-index-based alternatives, while maintaining the br-index’s favorable memory characteristics. For example, all available complete *E. coli* genomes on NCBI’s RefSeq collection can be compiled into a b-move index that fits into the RAM of a typical laptop. Thus, b-move proves practical and scalable for pan-genome indexing and querying. We provide a C++ implementation of b-move, supporting efficient lossless approximate pattern matching including locate functionality, available at https://github.com/biointec/b-move under the AGPL-3.0 license.

**Funding:** *Lore Depuydt*: PhD Fellowship FR (1117322N), Research Foundation – Flanders (FWO)

*Luca Renders*: PhD Fellowship SB (1SE7822N), Research Foundation – Flanders (FWO)

*Travis Gagie*: NSERC Discovery Grant RGPIN-07185-2020 to Travis Gagie and NIH grant R01HG011392 to Ben Langmead

## 1 Introduction

Since the advent of long-read sequencing platforms, the availability of high-quality genome sequences has increased dramatically. To exploit this data, it is becoming increasingly common to compile individuals from the same species or several related species into a single index, forming what is known as a pan-genome [10]. This approach aims to better capture genetic diversity and mitigate biased results stemming from the choice of reference.

Many widely used bioinformatics tools, such as BWA [19] and Bowtie 2 [18], rely on the FM-index [12]. The FM-index, based on the Burrows-Wheeler transform (BWT) [7] and suffix array (SA) [21], efficiently counts and locates exact occurrences of a search pattern in the reference. While it is compact and fast for relatively small references (e.g., a single human reference genome or a few dozen bacterial species), its memory use scales linearly with the total genome content. This limitation calls for new index types capable of storing and analyzing large genome collections within the memory constraints of modern workstations and laptops.

The BWT’s inherent compressibility (see e.g., bzip2 [27]), particularly for highly repetitive input texts like pan-genomes, has led to a focus on run-length compression. The run-length FM-index (RLFM-index) [20] efficiently counts occurrences in *O*(*r*) space, *r* being the number of character runs in the BWT, but requires *O*(*n*) additional space for locating functionality. Recently, Gagie et al. [13, 14] introduced the r-index, which also supports locating functionality in *O*(*r*) space. Its reduced memory requirements have made the r-index the foundation for several tools, including MONI [26], PHONI [5], SPUMONI [1], and SPUMONI 2 [2]. Nishimoto and Tabei [22] more recently proposed the ‘move structure’, a run-length compressed index achieving, unlike the r-index, both *O*(*r*) space and *O*(1) LF operations. Built upon this, Movi [28] offers efficient pan-genome index building and querying, matching SPUMONI’s functionality but with significantly faster performance.

Despite these advancements, a notable limitation persists: aforementioned indexes and tools support only backward stepping through the LF operation, limiting the range of queries possible. Specifically, tools relying on the r-index or move structure focus almost exclusively on MEM-finding, using (pseudo-)matching lengths and statistics. In a pan-genome with thousands of genomes, this approach could yield an overwhelming number of MEMs, making downstream full read alignment based on seed-and-extend algorithms challenging and potentially infeasible [4].

Recognizing this limitation, Arakawa et al. [3] introduced the br-index. This extension of the traditional r-index enables bidirectional match extensions during the search process, i.e., both to the left and right, in arbitrary order. While the br-index offers more operational flexibility, supporting functionalities like approximate pattern matching (APM) based on the pigeonhole principle or more general search schemes [16], it also inherits the high computational overhead of the r-index. This overhead stems from the intricate interplay of rank and select queries on compressed sparse bitvectors and wavelet trees, leading to multiple cache misses. Despite its favorable *O*(*r*) memory complexity, the br-index can be one order of magnitude slower than the bidirectional FM-index, hindering its adoption in practical bioinformatics tools.

### Contribution

In this paper, we introduce b-move, a bidirectional extension of the move structure, as a faster alternative to the br-index. This paper is organized as follows. In Section 2, we recapitulate basic concepts and existing methods that form the foundation of this paper. In Section 3, we propose our bidirectional move structure, with a detailed description of its core bidirectional character extension functionalities. In Section 4, we demonstrate its efficiency in performing synchronized bidirectional character extensions, achieving a speedup of 6 to 8 times compared to the br-index and showing performance comparable to FM-index-based tools. We confirm b-move’s favorable *O*(*r*) memory complexity, superior to the bidirectional FM-index both in theory and in practice.

## 2 Preliminaries

In this paper, arrays and strings are indexed starting from zero. Consider a search text *T* with a length of *n* = |*T* | over an alphabet ∑. In the context of pan-genomes, *T* consists of multiple concatenated genome sequences. We assume that *T* ends with the sentinel character $, which is lexicographically smaller than any other character in ∑. A substring within string *T* is represented as an interval [*i, j*] over *T*, where 0 ≤ *i* ≤ *j* ≤ *n* − 1. The *i*th suffix of *T*, denoted as *T*_*i*_, refers to the substring *T* [*i, n* − 1].

In this paper, we primarily focus on accelerating bidirectional character extensions, which are based on the LF operation. These extensions are fundamental to the counting functionality, which returns the number of occurrences of a specific pattern in the search text. Consequently, our discussion will be limited to the data structures required for these character extensions and the computations needed to perform them.

### 2.1 Counting exact occurrences with the FM-index and r-index

The uncompressed FM-index supports counting functionality in *O*(*n*) space and *O*(*m*) time, where *m* is the length of pattern *P* [12]. Consider the interval [*s, e*] in the FM-index corresponding to all sorted suffixes of *T* prefixed by pattern *P*. The interval [*s*^*′*^, *e*^*′*^] for *P* ‘s extension *cP* can be found as *s*^*′*^ = C[*c*]+rank_*c*_(BWT, *s*) and *e*^*′*^ = C[*c*]+rank_*c*_(BWT, *e*+1)−1. Here, C[*c*] denotes the number of characters in *T* strictly smaller than *c*, and rank_*c*_(BWT, *i*) counts the occurrences of character *c* in the BWT before index *i*. These operations can be executed in constant time if the BWT is represented as a collection of |∑| bitvectors with constant-time rank support. For readers less familiar with counting functionality using backward search in the FM-index, a more extensive overview is provided in [11].

To address the FM-index’s space inefficiency, which increases linearly with the size of the search text, Gagie et al. [13, 14] introduced the r-index. The r-index offers counting functionality in *O*(*r*) space with a time complexity of *O*(*m* · log(log_*w*_(|∑| + *n/r*))), where *w* = Ω(log(*n*)) is the machine word size. Although the counting process in the r-index is conceptually similar to that in the FM-index, it requires a more complex combination of operations. Specifically, each character extension involves a combination of access, rank, select, and conditional operations performed on multiple data structures, which, collectively, cannot be performed in constant time. For further details, refer to [14]. This complexity leads to more random access operations and, consequently, cache misses.

### 2.2 Constant-time LF operations with the move structure

The LF(*i*) operation maps the character at index *i* in the BWT or *L* (the last column of the lexicographically ordered rotations of *T*) to its corresponding character in *F* (the first column of the lexicographically ordered rotations of *T*). Nishimoto and Tabei [22] introduced the move structure as an alternative to the r-index that achieves LF operations in constant time. Their key insight is that LF mappings within a single run map to consecutive positions in *F*. Instead of mapping a single position in *L* to *F*, *input intervals* in *L* are mapped to *output intervals* in *F*. For example, in the index illustrated in Table 1, the input interval [2, 5] from the second BWT run corresponds to the output interval [11, 14] in *F* .

**Table 1.**
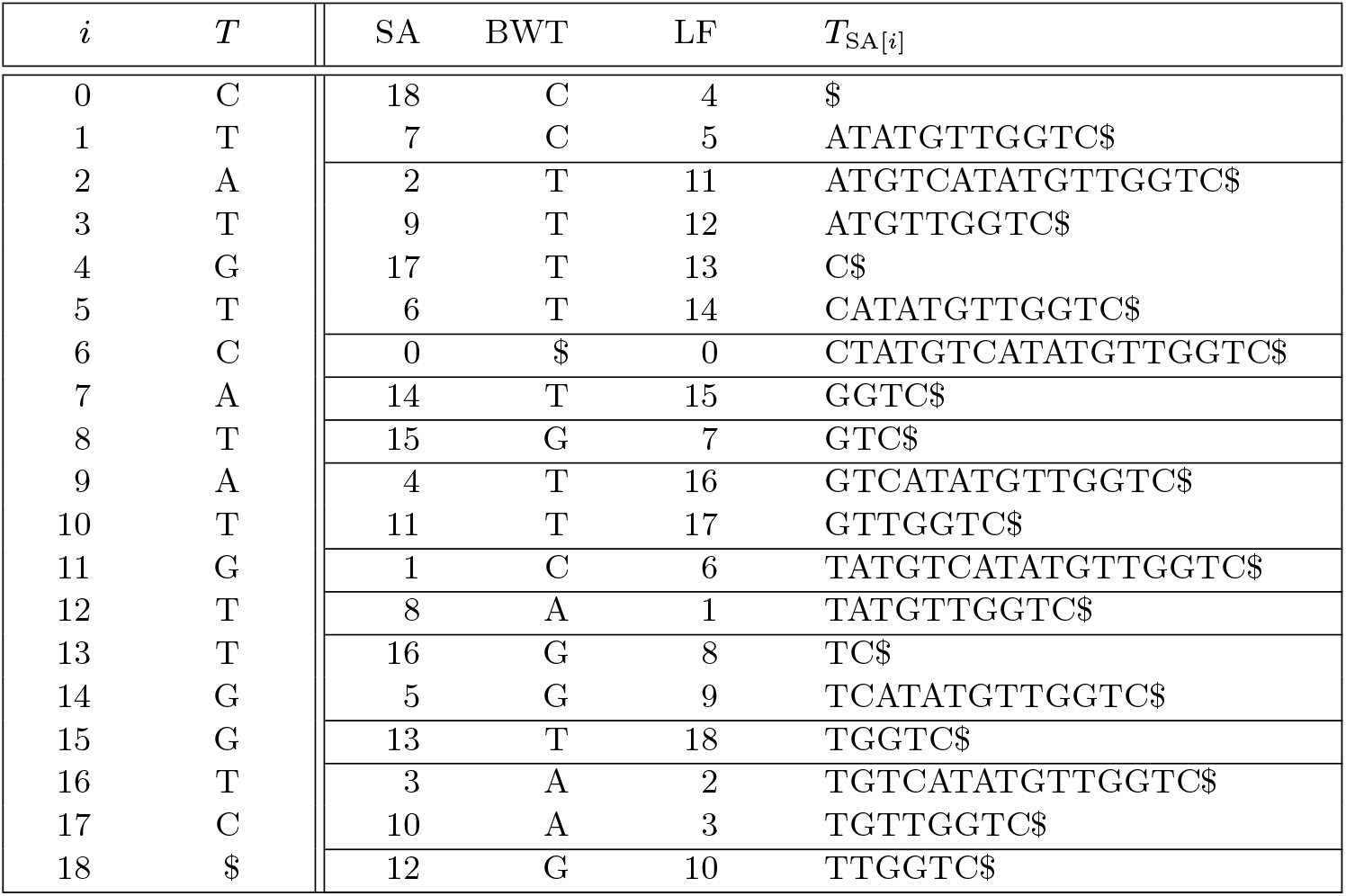
Search text *T* = “CTATGTCATATGTTGGTC$” with its suffix array SA, Burrows-Wheeler transform BWT or *L*, LF mapping, and suffixes (the first characters of which represent *F*). BWT runs are delineated by horizontal lines.

#### Algorithm 1

Find LF(*i*) given tuple (*i, j*), where *j* is the run index that contains BWT index *i*, using the move structure *M* .

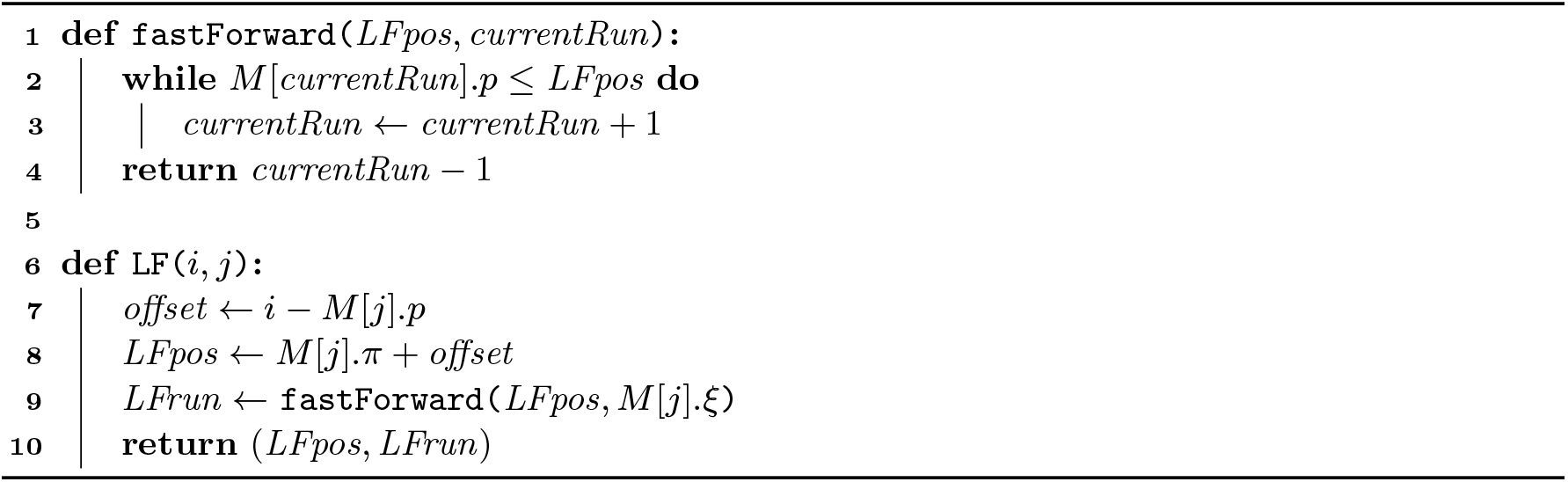

Conceptually, the move structure is a table with *r* rows: one for each run in the BWT. Following the notation from Zakeri et al. [28], each row contains four elements: the character *c* of the run, the starting index *p* of its input interval, the starting index *π* = LF(*p*) of its output interval, and the run index *ξ* that contains *π*. The move structure *M* for the example of Table 1 is shown in Table 2. Note that *ξ* is not injective: multiple input runs can have their *π* residing in the same output run.

**Table 2.**
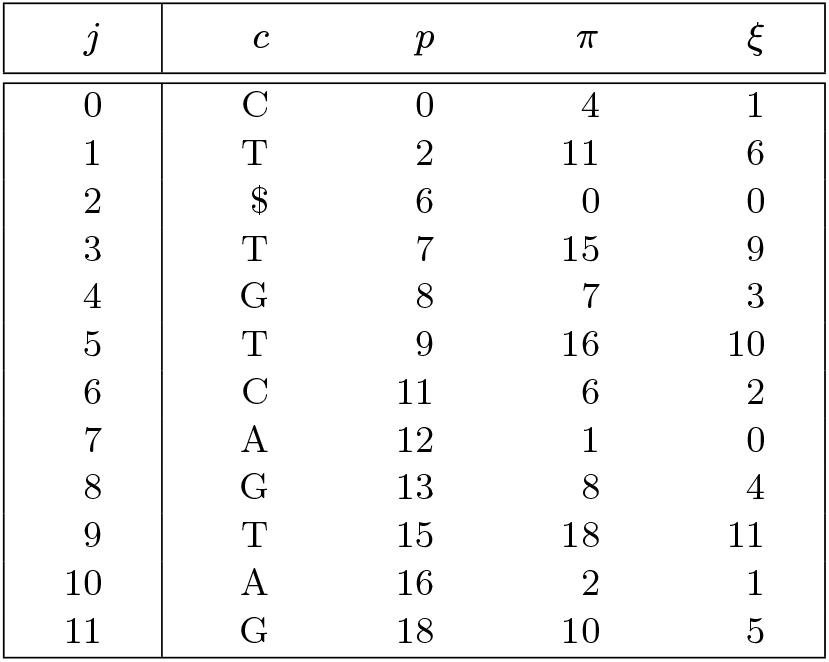
Move structure *M* corresponding to search text *T* = “CTATGTCATATGTTGGTC$”.

| $j$ | $c$ | $p$ | $\pi$ | $\xi$ |
| --- | --- | --- | --- | --- |
| 0 | C | 0 | 4 | 1 |
| 1 | T | 2 | 11 | 6 |
| 2 | \$ | 6 | 0 | 0 |
| 3 | T | 7 | 15 | 9 |
| 4 | G | 8 | 7 | 3 |
| 5 | T | 9 | 16 | 10 |
| 6 | C | 11 | 6 | 2 |
| 7 | A | 12 | 1 | 0 |
| 8 | G | 13 | 8 | 4 |
| 9 | T | 15 | 18 | 11 |
| 10 | A | 16 | 2 | 1 |
| 11 | G | 18 | 10 | 5 |

To perform the LF operation on a given BWT index *i*, we need the run index *j* that contains *i* to access move structure *M*. Assuming we know both *i* and *j* and aim to compute LF(*i*), Algorithm 1 outlines the process. We start by determining the offset between *i* and the start of its run. With this offset and the start of the output interval for run *j*, we compute LF(*i*). For potential subsequent LF operations, we must also find the run index containing LF(*i*) using fast forwarding functionality. Fast forwarding is necessary when run *M* [*j*].*ξ* does not contain LF(*i*). For example, to find the run index corresponding to LF(5) = 14, where *i* = 5 resides within BWT run *j* = 1, we observe that run index *M* [1].*ξ* = 6 (covering the interval [11, 11]) does not include BWT index 14. In such cases, we traverse subsequent runs until the correct run is located using the `fastForward` function. We ensure that the access on line 2 is never out-of-bounds by adding an extra row at the bottom of the move structure *M*, such that *M* [*r*].*p* = *n*.

The LF operation, as described in Algorithm 1, is more cache-friendly than the alternative in the r-index. It involves jumping to row *M* [*M* [*j*].*ξ*] and possibly accessing subsequent rows. Fetching these subsequent rows translates to linear memory access (streaming) and can be efficiently performed in contemporary computer architectures. For true constant-time LF operations, the number of steps in the fast forwarding function should be limited. While Nishimoto and Tabei [22] suggested balancing or splitting the input and output intervals to achieve this, Zakeri et al. [28] found that splitting these intervals did not result in a notable speedup when implementing Movi. Therefore, we maintain the original runs in this paper. Technically, this leads to an *O*(*r*) worst-case time complexity. In practice, however, this is the more efficient choice for building and storing the index and does not noticeably impact search performance.

The move structure principle can also provide for constant-time *ϕ* and *ϕ*^*−*1^ operations. However, in this paper, we focus solely on accelerating character extensions, for which this optimization is not applicable.

### 2.3 Finding approximate occurrences with the bidirectional FM-index and br-index

The bidirectional FM-index [17] can extend a pattern *P* to *Pc* or *cP* by maintaining synchronized intervals over SA and SA^rev^. Here, SA^rev^ is the suffix array of *T* ^rev^ (the reverse of *T*). To enable bidirectional extension, certain components corresponding to the FM-index of *T* ^rev^ are stored; see [11] for details. Consider intervals [*s, e*] and [*s*^rev^, *e*^rev^] for *P* in SA and *P* ^rev^ in SA^rev^. To extend *P* to *cP*, we find [*s*^*′*^, *e*^*′*^] for *cP* in SA as in Section 2.1. Updating [*s*^rev^, *e*^rev^] involves recognizing that [*s*^rev*′*^, *e*^rev*′*^] ⊆ [*s*^rev^, *e*^rev^] due to *P* ^rev^*c* being prefixed by *P* ^rev^. Moreover, all suffixes in SA^rev^ prefixed by *P* ^rev^*a, a* ≺ *c*, are sorted before those within the interval [*s*^rev*′*^, *e*^rev*′*^]. If we compute the cumulative number of occurrences *x* of *aP* in *T*, for all characters *a* ≺ *c*, using the procedure from Section 2.1, interval [*s*^rev*′*^, *e*^rev*′*^] in SA^rev^ can be found as *s*^rev*′*^ = *s*^rev^ + *x* and *e*^rev*′*^ = *s*^rev^ + *x* + *y* − 1, where *y* = *e*^*′*^ − *s*^*′*^ + 1 is the count of *cP* occurrences in *T*. Extending *P* to *Pc* is done symmetrically.

Arakawa et al. [3] introduced the br-index, an extension of the r-index taking up *O*(*r*+*r*^rev^) space, *r*^rev^ being the number of runs in the BWT of *T* ^rev^. Interval synchronization in the br-index parallels the method above but uses more intricate combinations of access, rank, select, and conditional operations on various data structures. Similar to the r-index, this leads to multiple cache misses and hence, slower performance. Additionally, the problem exacerbates as the number of occurrences for all *a* ≺ *c* must be found, requiring *O*(|∑|) rank operations on BWT or BWT^rev^.

If locating the occurrences is necessary, a toehold update follows each character extension. This is also achieved with a combination of core operations like LF. For a more detailed overview, refer to [3].

Note that being able to match a pattern *P* in both directions, starting at any position within *P* and switching direction at any time, increases operational flexibility. For example, approximate matches of a pattern can be sought using the pigeonhole principle or, more generally, search schemes [16]. The demonstrated efficiency of search schemes, for example in lossless read-mapper Columba [25, 23, 24], motivates the need for fast bidirectional character extensions.

## 3 Bidirectional move structure

While Nishimoto and Tabei’s move structure with constant-time LF support shows promise, practical implementations remain scarce. Brown et al. [6] analyzed time-space tradeoffs and proposed a compressed implementation of the move table that efficiently counts exact occurrences while using less space. Zakeri et al. [28] introduced Movi, a fast and cache-efficient full-text pan-genome index that supports computing matching statistics and pseudo-matching lengths, useful for finding maximal exact matches. However, to our knowledge, no practical bidirectional move structure exists that supports finding approximate occurrences of a pattern in a search text. In this section, we introduce our bidirectional move structure, which (cache-)efficiently performs bidirectional character extensions using *O*(*r* + *r*^rev^) space.

### 3.1 Backward search

We elaborate on unidirectional search based on the LF functionality discussed in Section 2.2. As for the LF operation, run indices corresponding to the current SA interval boundaries are required for character extension. Consider interval [*s, e*] in SA for pattern *P*, with *R*_*s*_ and *R*_*e*_ the runs containing *s* and *e*. To extend *P* to *cP*, the LF functionality cannot be applied directly to the interval boundaries, as runs *R*_*s*_ and *R*_*e*_ may not match *c*. Instead, LF must be performed on subinterval [*s*_*c*_, *e*_*c*_] ⊆ [*s, e*], which is the smallest interval containing all occurrences of *c* in the BWT in [*s, e*]. We again require run indices *R*_*s*,*c*_ and *R*_*e*,*c*_ for *s*_*c*_ and *e*_*c*_. To find *R*_*s*,*c*_ and *R*_*e*,*c*_ using only the move structure, we simply walk down and up along the rows until finding an instance of *c*, inspired by Zakeri et al.’s repositioning in Movi-default [28]. Function `walkToNextRun` in Algorithm 2 demonstrates this approach. If no occurrences of *c* exist within [*s, e*] (checked on line 7), −1 is returned for both values.

#### Algorithm 2

Let [*s, e*] be the interval over SA corresponding to *P*, which we want to extend to *cP*. *R*_*s*_ and *R*_*e*_ are the runs containing *s* and *e*, respectively. Functions `walkToNextRun` and `walkToPreviousRun` return the SA indices *s*_*c*_ and *e*_*c*_ (along with their run indices *R*_*s*,*c*_ and *R*_*e*,*c*_) which indicate the smallest subinterval within [*s, e*] containing all occurrences of *c* in the BWT.

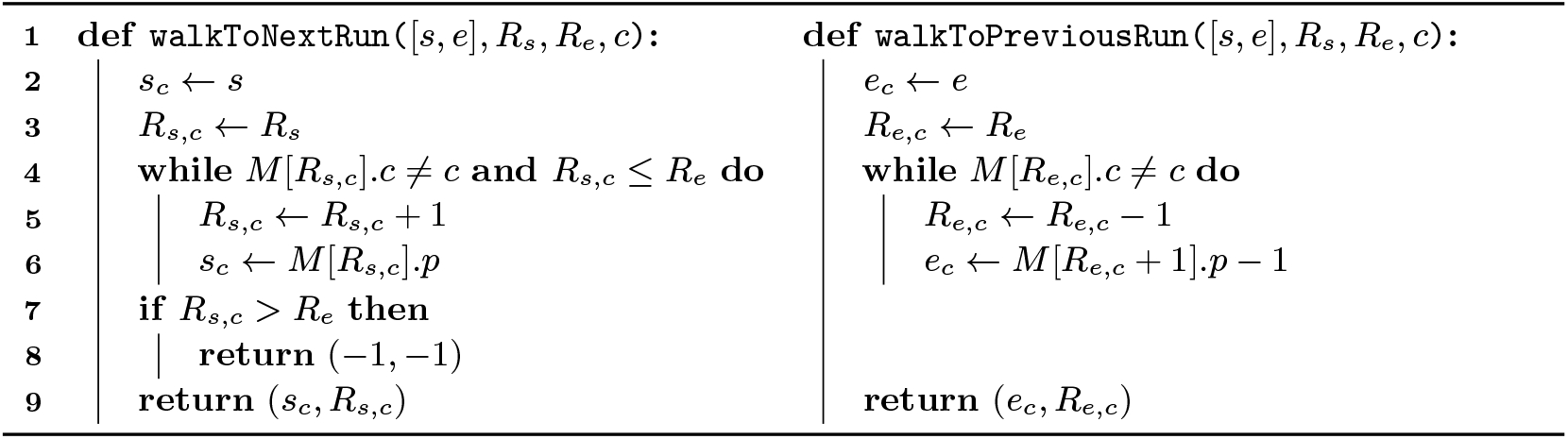

#### Algorithm 3

Extend pattern *P*, represented by [*s, e*], corresponding to respectively run index *R*_*s*_ and *R*_*e*_, to *cP*. The algorithm returns the updated interval [*s*^*′*^, *e*^*′*^], corresponding to respectively run index *R*^*′*^ and *R*^*′*^, for *cP* .

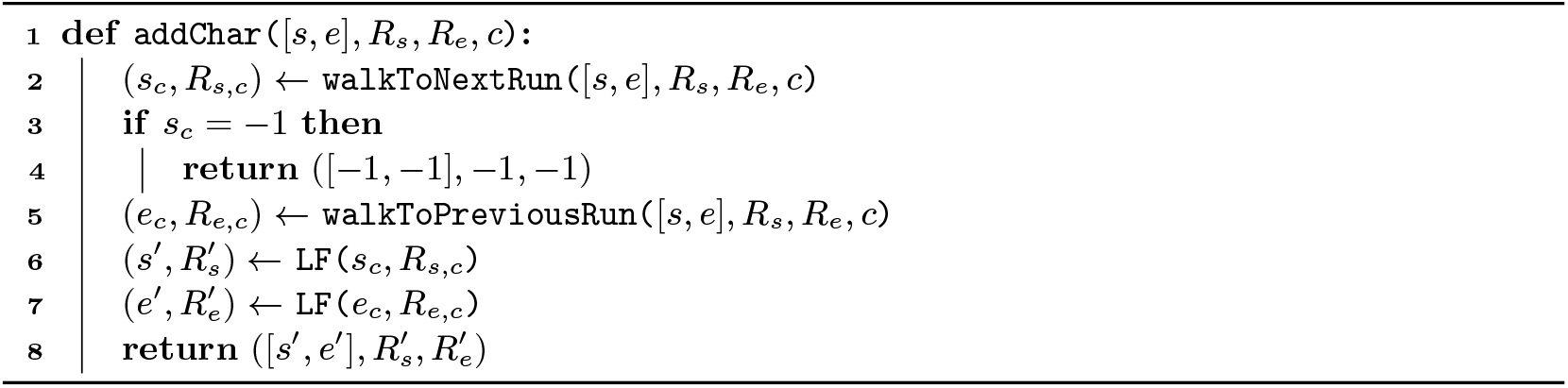

Function `walkToPreviousRun` in Algorithm 2 is similar to function `walkToNextRun`, but the check on line 7 is no longer necessary as the function is always executed in circumstances where it is known that *c* occurs in the BWT within [*s, e*] (usually by running walkToNextRun first). Also, on line 6 in function `walkToPreviousRun`, a subsequent row is accessed since the end indices of the runs are not stored in the move table. Note that this access can never be out-of-bounds. Function `walkToPreviousRun` can also be used to update the toehold if required for locating, details are omitted here.

Algorithm 3 then combines the walking from Algorithm 2 and the LF from Algorithm 1 into the functionality to extend a pattern *P* to *cP*. Algorithms 2 and 3 have a worst-case time complexity of *O*(*r*) but are very fast in practice. We also explored constant-time alternatives for Algorithm 2, such as storing additional bitvectors representing the run heads, supporting rank and select operations. However, this option performed worse in practice, both in memory usage and speed, due to complex memory access patterns.

### 3.2 Bidirectional search

Similar to the bidirectional FM-index and r-index, we incorporate an additional component to represent the reverse search text *T* ^rev^ for bidirectional search with our bidirectional move structure. Specifically, we store the move table *M* ^rev^ corresponding to *T* ^rev^. Table 3 shows *M* ^rev^ corresponding to *M* in Table 2. For completeness, the FM-index corresponding to *T* ^rev^ is shown in Table S1.

**Table 3.** Move structure *M* ^rev^ corresponding to search text *T* ^rev^ = “$CTGGTTGTATACT-GTATC”.

| $j$ | $c$ | $p$ | $\pi$ | $\xi$ |
| --- | --- | --- | --- | --- |
| 0 | C | 0 | 4 | 1 |
| 1 | T | 1 | 11 | 6 |
| 2 | \$ | 5 | 0 | 0 |
| 3 | A | 6 | 1 | 1 |
| 4 | T | 7 | 15 | 9 |
| 5 | G | 10 | 7 | 4 |
| 6 | A | 11 | 2 | 1 |
| 7 | G | 12 | 8 | 4 |
| 8 | A | 14 | 3 | 1 |
| 9 | C | 15 | 5 | 2 |
| 10 | T | 16 | 18 | 12 |
| 11 | C | 17 | 6 | 3 |
| 12 | G | 18 | 10 | 5 |

Using the combination of *M* and *M* ^rev^, we can synchronize the intervals corresponding to a search pattern *P* over SA and *P* ^rev^ over SA^rev^. Algorithm 4 details how pattern *P* can be extended to *cP* while keeping both intervals up to date. The approach is conceptually similar to that described in Section 2.3. Extending *P* to the right, i.e., to *Pc*, is analogous and is detailed in supplementary Algorithm S1.

#### Memoization

For clarity, the algorithms were built step-by-step. Thus, the time complexity for Algorithms 4 and S1 is *O*(|∑| · *r*) and *O*(|∑| · *r*^rev^), respectively, as all characters *a* ≺ *c* must be checked (line 8). Alternatively, combining Algorithms 1 to 4 can achieve *O*(*r*) bidirectional character extensions instead of *O*(|∑| · *r*). As |∑| is small in the context of this paper, this is mostly a theoretical improvement. To achieve this, consider the calls to function `addChar` in the for-loop on line 8 (and on line 2) of Algorithm 4, causing the |∑| factor. As Algorithm 3 is called |∑| times, the following *O*(*r*) time functions are also executed |∑| times:

i. `walkToNextRun` (Alg. 2)
ii. `walkToPreviousRun` (Alg. 2)
iii. `LF` (Alg. 1), twice

For `walkToNextRun` (i), the while-loop on line 4 in Algorithm 2 could be merged with the for-loop on line 8 of Algorithm 4 to continue walking until the first occurrence of each character *a* ⪯ *c* is found. This results in one *O*(*r*) walk, keeping each (*s*_*a*_, *R*_*s*,*a*_) tuple in memory. Similarly, `walkToPreviousRun` (ii) can be reduced to one *O*(*r*) walk, keeping each (*e*_*a*_, *R*_*e*,*a*_) tuple in memory.

Then, on each (*s*_*a*_, *R*_*s*,*a*_) and (*e*_*a*_, *R*_*e*,*a*_) tuple, `LF` (iii) is performed, which is *O*(*r*) due to the `fastForward` procedure. However, fast forwarding is always limited to the output interval corresponding to the input run/interval. Since the input runs for different characters *a* ⪯ *c* are distinct, fast forwarding occurs in disjoint output intervals, collectively summing to *O*(*r*) steps. If *R*_*s*,*a*_ = *R*_*e*,*a*_ (i.e., they have the same output interval), redundant fast forwarding can be avoided through memoization as well.

##### Algorithm 4

Update intervals ([*s, e*], *R*_*s*_, *R*_*e*_) and ([*s*^rev^, *e*^rev^],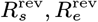) corresponding to *P*, to intervals ([*s*^*′*^, *e*^*′*^], *R*^*′ s*^, *R*^*′ s*^) and ([*s*^rev*′*^, *e*^rev*′*^],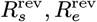) that correspond to *cP* .

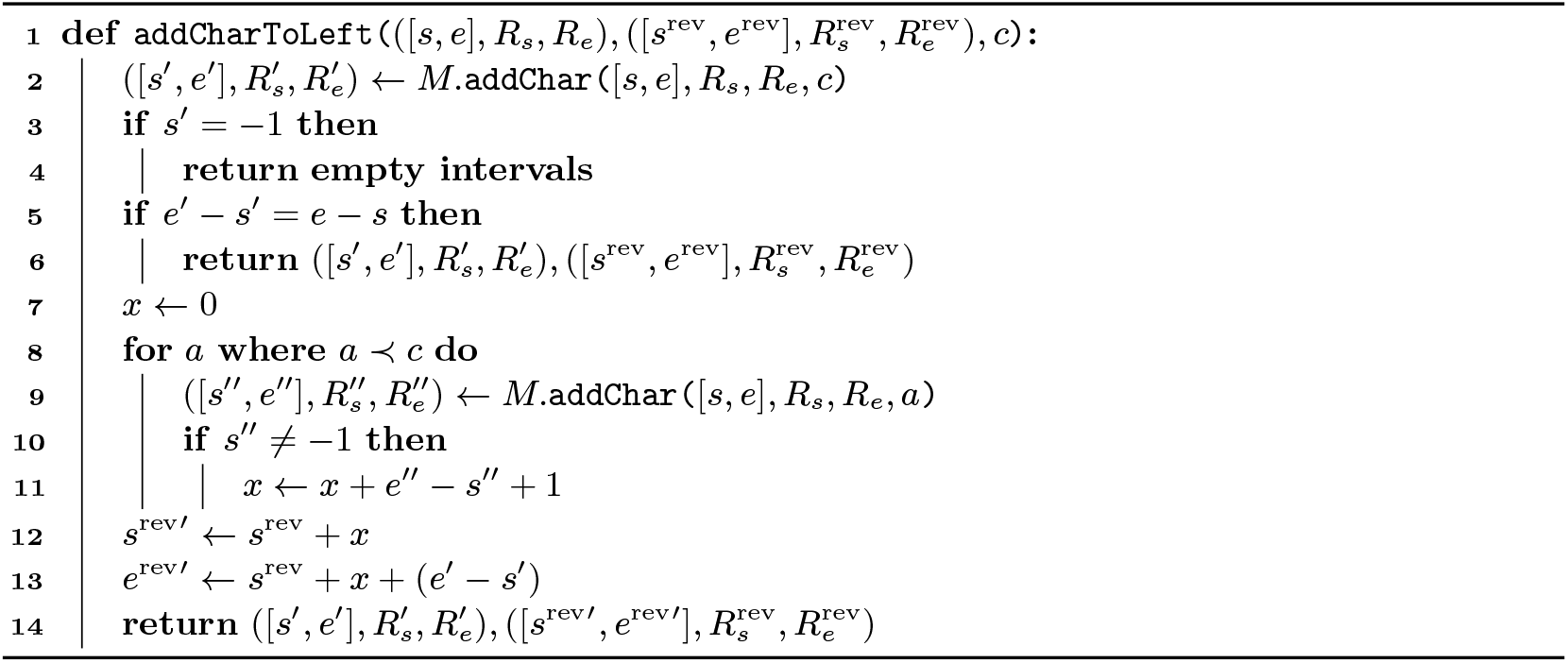

##### Algorithm 5

Update run indices using binary search to find the correct run containing index *i*. The algorithm uses the move table *M* upon which it is called.

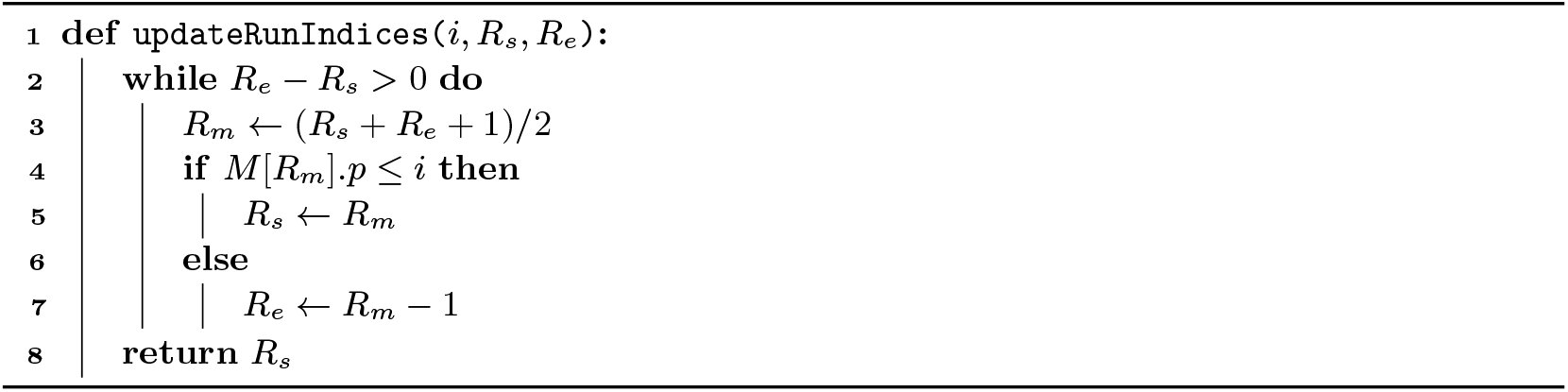

As such, the total combination of walking steps and fast forwarding steps necessary to perform bidirectional character extensions sums to *O*(*r*) instead of *O*(|∑| · *r*). Analogously, Algorithm S1 can be performed in *O*(*r*^rev^) time. We emphasize that in practice, the number of walking/fast forwarding steps is small, and due to linear memory access (streaming), the bidirectional character extensions are very fast.

### 3.3 Direction switches

In Algorithm 4, run indices 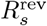 and 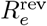 are not updated, and the same holds for *R*_*s*_ and *R*_*e*_ in Algorithm S1. Updating these indices at each synchronization step is not efficient nor necessary; they are required only when switching search direction.

Consider pattern *P* =“TATGTTGGT”, split into two parts: “TATGT|TGGT”. For the sake of the example, we match the first part to the left starting at the separation point. After five calls to `addCharToLeft`, the state is ([*s, e*], *R*_*s*_, *R*_*e*_), ([*s*^rev^, *e*^rev^], 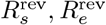) = ([11 12] 6 7) ([16 17] 0 12). Indeed, neither SA^rev^ index 16 nor 17 lies within their corresponding runs of 0 and 12. To switch direction and extend the match to the right, we first update 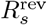 and 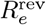.

We employ a binary search over the move rows, starting with the current (outdated) run indices, detailed in Algorithm 5. For our example, *M* ^rev^.`updateRunIndices` (16, 0, 12) = 10 and *M* ^rev^.`updateRunIndices` (17, 0, 12) = 11. With the correct run indices, we can extend the match to the right.

Algorithm 5 has a time complexity of *O*(log(*r*)). Note that in practice, direction switches are infrequent. Moreover, the binary search often operates with narrow starting intervals. In the example above, we started from the complete interval over all move rows. For another switch back to the left, we would call *M*. `updateRunIndices` (*i*, 6, 7).

## 4 Results

### 4.1 Data and hardware

We built pan-genomes from two datasets^1^: i) 512 human chromosome 19 haplotypes from the 1000 Genomes Project [9], and ii) 3264 *Escherichia coli* strains downloaded from NCBI’s Reference Sequence (RefSeq) collection. Characters ‘N’ were removed from these genomes. The chromosomes or strains are concatenated into one string, from which the indexes are built. Table 4 provides more detailed statistics regarding the pan-genomes we used for benchmarking. Note that the last column with the ratio between *n* and *r* confirms that the human pan-genome is more repetitive and conserved than the bacterial pan-genome.

**Table 4.** Details of the pan-genomes that are used for benchmarking. For each pan-genome, we report the number of chromosomes or strains it contains, the length *n* of the corresponding concatenated string *T*, the number of runs *r* in the BWT of *T*, the number of runs *r*^rev^ in the BWT of *T* ^rev^, and the ratio of *n* and *r*.

| Dataset | # | $n/10^6$ | $r/10^6$ | $r^{\text{rev}}/10^6$ | $n/r$ |
| --- | --- | --- | --- | --- | --- |
| Human chrom. 19 | 2 | 111.61 | 31.16 | 31.16 | 3.58 |
|  | 4 | 223.22 | 31.63 | 31.63 | 7.06 |
|  | 8 | 446.44 | 32.00 | 32.00 | 13.95 |
|  | 16 | 892.89 | 32.40 | 32.40 | 27.56 |
|  | 32 | 1 785.77 | 32.83 | 32.82 | 54.40 |
|  | 64 | 3 571.53 | 33.34 | 33.34 | 107.12 |
|  | 128 | 7 143.04 | 34.05 | 34.05 | 209.76 |
|  | 256 | 14 286.12 | 35.62 | 35.62 | 401.07 |
|  | 512 | 28 572.24 | 39.24 | 39.24 | 728.22 |
| *E. coli* | 2 | 10.08 | 5.06 | 5.06 | 1.99 |
|  | 4 | 20.11 | 9.11 | 9.11 | 2.21 |
|  | 8 | 40.16 | 11.43 | 11.43 | 3.51 |
|  | 16 | 80.14 | 15.17 | 15.17 | 5.28 |
|  | 32 | 158.30 | 19.42 | 19.42 | 8.15 |
|  | 64 | 317.21 | 27.34 | 27.35 | 11.60 |
|  | 128 | 630.15 | 34.68 | 34.68 | 18.17 |
|  | 256 | 1 262.28 | 52.06 | 52.06 | 24.24 |
|  | 512 | 2 527.97 | 64.20 | 64.20 | 39.38 |
|  | 1 024 | 5 075.37 | 78.79 | 78.79 | 64.42 |
|  | 2 048 | 10 172.03 | 98.74 | 98.74 | 103.02 |
|  | 3 264 | 16 207.62 | 118.15 | 118.15 | 137.18 |

For benchmarking the human pan-genomes, we consider 100 000 Illumina HiSeq 2500 reads (151 bp) randomly sampled from a larger whole genome sequencing dataset (accession number SRR17981962). For the bacterial pan-genomes, we use 100 000 Illumina NovaSeq 6000 reads (151 bp) randomly sampled from a larger whole genome sequencing dataset (accession number SRR28249370). Benchmarks were performed on a Red Hat Enterprise Linux 8 system, using a single core of two 18-core Intel^®^ Xeon^®^ Gold 6140 CPUs running at a base clock frequency of 2.30 GHz with 177 GiB of RAM. To account for variability in runtime, all reported runtimes are averaged over 10 runs. They exclude the time to read the index from and write the output to disk.

### 4.2 Character extension performance

In Figure 1, we compare the efficiency of bidirectional character extensions using the original br-index implementation^2^ by Arakawa et al. [3], and our bidirectional move structure, referred to as b-move. The left panel displays the total time for finding all SA intervals for occurrences up to a specific number of mismatches (i.e., no *ϕ* operations). To ensure a fair comparison, we adjusted b-move’s parameters to closely match the implementation of the br-index (e.g., Hamming distance, the use of the pigeonhole principle, and no further optimizations). As shown in the chart, b-move outperforms the br-index by a factor of 5 to 8 in terms of total search time, with the speed-up increasing with the allowed number of mismatches.

**Figure 1.**
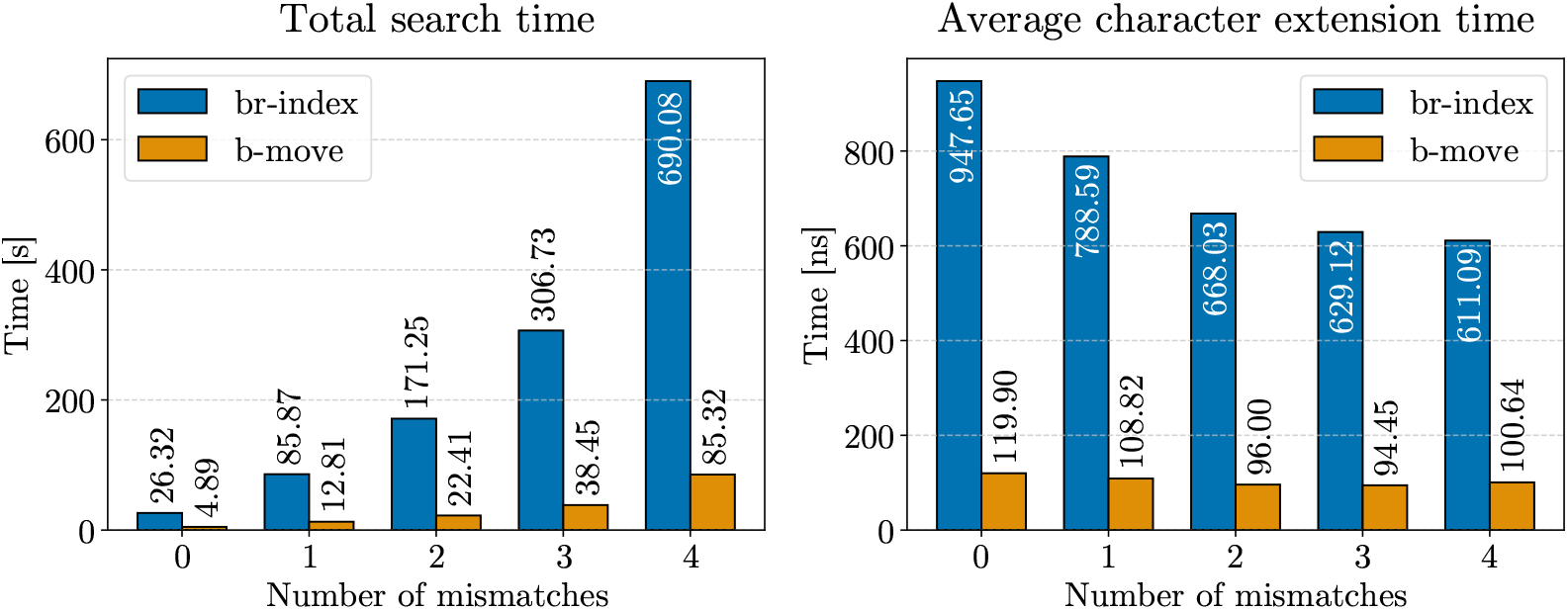
Benchmark results for finding all SA intervals corresponding to all occurrences up to a certain number of mismatches (x-axis). The left panel displays the total search time, while the right panel shows the average execution time for the core bidirectional character extension functionality. We aligned 100 000 Illumina reads of length 151 bp and their reverse complements to the pan-genome composed of 512 human chromosome 19 haplotypes.

While we aimed for similar parameters, implementation differences beyond core character extension functionality may affect the total search time. To address this, we specifically benchmarked the character extension functionality described in Algorithms 4 and S1, along with their counterparts in the br-index. We used a built-in instruction (constant rdtscp) to count CPU cycles and scaled the average cycle count to time using the clock frequency. Given that cache misses consume dozens of CPU cycles, the time per character extension also serves as a proxy for cache misses. The right panel of Figure 1 shows a speed-up of 6 to 8 × in favor of b-move. The time for a single bidirectional character extension in the br-index decreases when allowing more mismatches. This is because with more allowed paths in the search tree, certain memory accesses are repeated more frequently, either to determine the widths of all intervals for *a* ≺ *c* or to extend with *a* itself. Consequently, the number of cache misses also decreases somewhat at higher error rates. This effect is mitigated in b-move due to its superior cache efficiency.

Additionally, we analyzed the *O*(*r*) and *O*(*r*^rev^) operations in each bidirectional character extension (see Section 3.2). Figure 2 shows the number of walking and fast forwarding steps for each successful bidirectional character extension from the same experiment as Figure 1 (3 mismatches). Most extensions require very few steps (note the log scale): 95% need fewer than 4 walking steps and fewer than 8 fast forwarding steps. This observation is confirmed by the median number of 0 walking steps and 1 fast forwarding step per extension (note that at least 1 fast forwarding step is required for each `LF` operation, see Alg. 1). Thus, these worst-case time complexities have minimal impact on actual performance.

**Figure 2.**
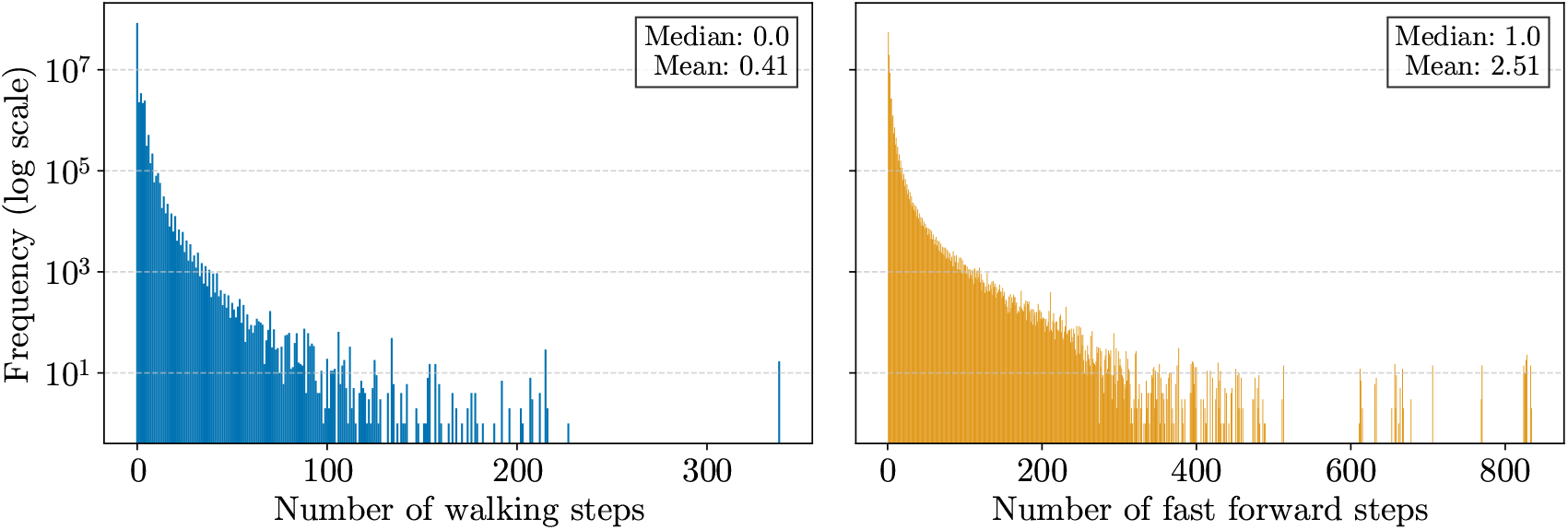
Log scale histograms for the number of walking and fast forwarding steps required per successful bidirectional character extension (96 363 328 in total) for finding all SA intervals corresponding to all occurrences up to 3 mismatches. We aligned 100 000 Illumina reads of length 151 bp and their reverse complements to the pan-genome composed of 512 human chromosome 19 haplotypes.

### 4.3 Complete approximate pattern matching performance

To leverage the efficient bidirectional character extensions from the previous section to their full potential, we extended b-move into a practical tool for lossless approximate pattern matching of reads to large pan-genomes. This means we report *all* occurrences within a pre-specified Hamming or edit distance. This functionality is similar to that provided by lossless read-mapper Columba (developed in the same research group) [25, 23, 24], but Columba’s index is based on the bidirectional FM-index (*O*(*n*) memory requirements). To ensure practical efficiency in b-move, we incorporated several optimizations originally developed for Columba: optimized edit distance to reduce redundancy, superior search schemes replacing pigeonhole methods, a lookup-table to bypass matching the first 10-mers, dynamic pattern partitioning, and bit-parallel pattern matching.

In Figure 3, we compare the br-index, b-move with these optimizations in place, and Columba^3^ in terms of peak memory usage (left panels) and approximate pattern matching performance (right panels) across various pan-genome sizes. This performance evaluation includes both the human chromosome 19 pan-genomes (top panels) and the *E. coli* pan-genomes (bottom panels). For a fair comparison, note that optimizations in Columba requiring the original text are disabled.

**Figure 3.**
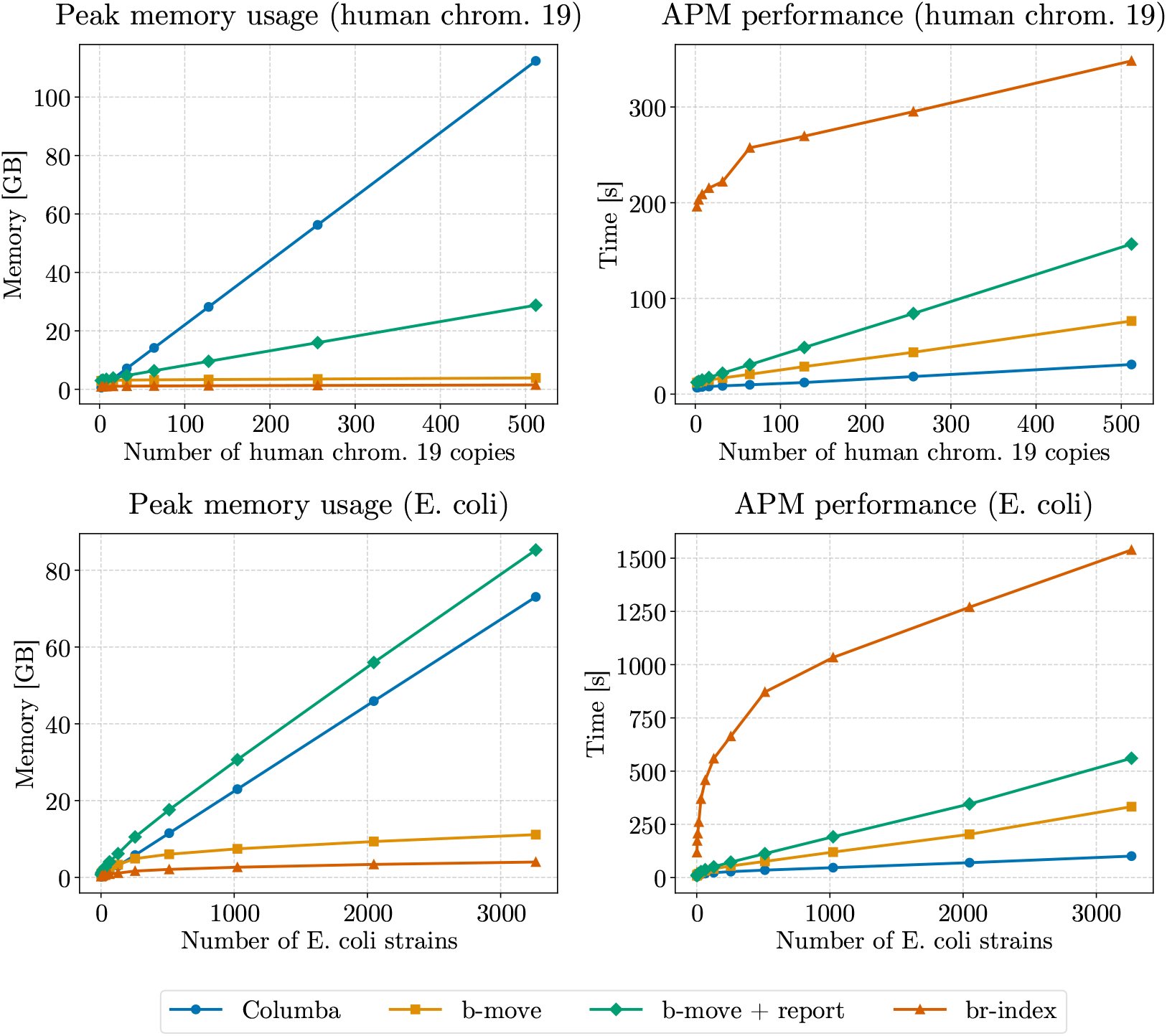
Benchmark results for approximate pattern matching using the br-index, b-move, and Columba (suffix array sparseness of 8). Additionally, we include results for b-move with reporting functionality, which is not included in the other results. We aligned 100 000 Illumina reads of length 151 bp and their reverse complements to pan-genomes for both human chromosome 19 and *E. coli*, across multiple sizes. We allowed for a maximum error distance of 3 (Hamming for br-index, edit for b-move and Columba).

Despite the fact that the *E. coli* pan-genomes are less repetitive than the human pan-genomes, both datasets demonstrate that the sublinear memory scaling characteristics of the br-index and b-move result in substantial memory reductions. Even for the largest pan-genomes, the br-index and b-move produce indexes that can stored in the RAM of a regular laptop. The difference in size between the br-index and b-move (more pronounced for *E. coli*) is put into perspective by the significant reduction relative to the FM-index-based tool. Regarding approximate pattern matching performance (including *ϕ*), b-move’s speed is of the same order of magnitude as Columba’s, albeit a constant factor of 2 or 3 larger. Conversely, the br-index is roughly 5 times slower than b-move.

In summary, b-move’s memory usage is orders of magnitude smaller than Columba’s for large, repetitive datasets, and closely resembles that of the br-index. Conversely, b-move outperforms the br-index by almost one order of magnitude while closely matching Columba’s performance in approximate pattern matching. Thus, we believe that b-move is the optimal index for scaling lossless approximate pattern matching to large pan-genomes.

Finally, the charts include a fourth line: b-move with reporting functionality. These results capture the overhead incurred by converting all found occurrences into SAM lines (no support for CIGAR strings yet) and maintaining them in memory throughout the search. Comparison between b-move + report and b-move alone highlights the substantial performance overhead of this reporting functionality. As a result, we excluded it from the main comparison. Nonetheless, an implementation for reporting is available both in b-move and Columba for users who require this feature. Note that the memory overhead due to reporting can be decreased significantly by writing out the alignments gradually throughout matching instead of buffering occurrences in memory (e.g., for *E. coli* the alignments comprise more space than Columba’s index). The performance overhead, however, is less straightforward to address. One option would be to explore different techniques for extracting the desired information without relying on locating, such as tagging the index with additional metadata [4].

### 4.4 Index size

Though the (bidirectional) move structure offers faster (see Section 4.2) character extensions than the (bidirectional) r-index with identical *O*(*r*) space complexity, its memory requirements are somewhat larger in practice. We evaluate this for the largest pan-genome (in terms of BWT runs) from Table 4: the pan-genome of 3 264 *E. coli* strains. A detailed overview of the space usage of the different components can be found in Table 5. For character extensions without locating functionality, the bidirectional move structure (5.9 GB) requires approximately 10 times more space than the bidirectional r-index (0.5 GB). Nevertheless, it is worth noting that our move structure, occupying less than 6 GB of memory, remains manageable even on a standard laptop.

**Table 5.** Detailed overview of the components in the bidirectional r-index and the bidirectional move structure for character extension support. The bottom section shows the additional memory required for supporting locating functionality, applicable to both previously discussed indexes.

| Component | Description | Memory usage for 3 264 *E. coli* genomes [MB] |
| --- | --- | --- |
| Bidirectional r-index as described in [3] |  |  |
| Reg. bv. runs | Sparse bitvector with $r$ 1-bits indicating run endings in the BWT of $T$ | 78.56 |
| Reg. bv. runs-per-letter | Division of the runs bitvector into $\vert \Sigma\vert $ shorter sparse bitvectors per character | 142.37 |
| Reg. run heads | Characters corresponding to runs in the BWT of $T$ | 49.13 |
| Rev. bv. runs | Analogous for $T^{\text{rev}}$ | 78.56 |
| Rev. bv. runs-per-letter | Analogous for $T^{\text{rev}}$ | 142.37 |
| Rev. run heads | Analogous for $T^{\text{rev}}$ | 49.13 |
| Total | - | 540.13 |
| Bidirectional move structure (this paper) |  |  |
| Reg. move table $M$ | Move table as described in Section 2.2 | 2 953.80 |
| Rev. move table $M^{\text{rev}}$ | Move table for $T^{\text{rev}}$ as described in Section 3.2 | 2 953.75 |
| Total | - | 5 907.55 |
| Extra data to support locate functionality [3] |  |  |
| Total | - | 3 306.51 |

For many bioinformatics applications that rely on read alignment, however, locating functionality is essential. When considering these additional data structures (detailed at the bottom of Table 5), the bidirectional move structure occupies 9.2 GB, compared to 3.8 GB required for the bidirectional r-index. In a broader context, this difference is reasonable given the substantial performance improvement offered by the move structure. This becomes even more evident when comparing the space usage with FM-index-based tools (as discussed in Section 4.3).

Note that the additional memory usage required to support locating can still be decreased by applying the subsampling technique [8, 15].

## 5 Conclusion

We propose b-move, a cache-efficient, run-length compressed index supporting lossless approximate pattern matching against large pan-genomes. b-move can perform character extensions 6 to 8 times faster than the br-index and is narrowing the performance gap with non-compressed FM-index-based pattern matching. Additionally, b-move demonstrates favorable sublinear memory characteristics, being orders of magnitude smaller than the FM-index for large pan-genomes. For example, *all* 3 264 available complete *E. coli* genomes on NCBI’s RefSeq collection can be compiled into a b-move index usable on a regular laptop. Future work includes investigating the application of the move principle to the *ϕ* and *ϕ*^*−*1^ operations to reduce cache misses in b-move’s locating functionality. Moreover, we aim to optimize the reporting functionality to minimize both memory (by altering the writing system) and runtime (by researching alternatives such as tagging) overhead. We also aim to integrate subsampling functionality to further reduce memory usage of the index itself. The source code of b-move is written in C++ and is available at https://github.com/biointec/b-move under AGPL-3.0 license.

## Acknowledgments

The authors thank Ben Langmead, Nathaniel Brown, and Mohsen Zakeri for their helpful feedback and suggestions.

---

**Table S1.**
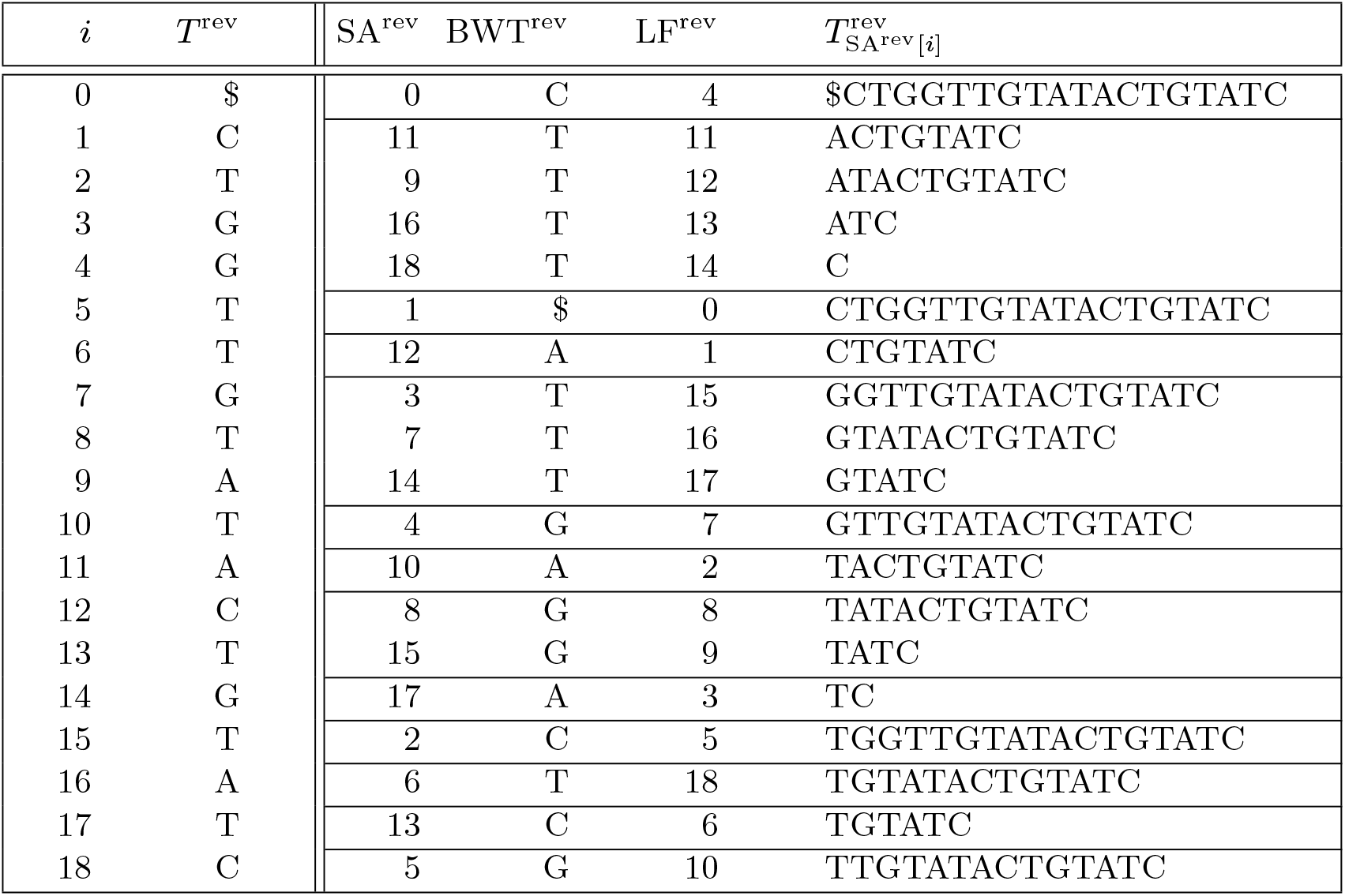
Reverse search text *T* ^rev^ = “$CTGGTTGTATACTGTATC” with its suffix array SA^rev^, Burrows-Wheeler transform BWT^rev^, LF mapping LF^rev^, and suffixes. BWT runs are delineated by horizontal lines.

## Algorithm S1

Update intervals ([*s, e*], *R*_*s*_, *R*_*e*_) and ([*s*^rev^, *e*^rev^], 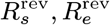) corresponding to *P*, to intervals ([*s*^*′*^, *e*^*′*^], *R*_*s*_, *R*_*e*_) and ([*s*^rev*′*^, *e*^rev*′*^],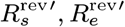) that correspond to *Pc*.

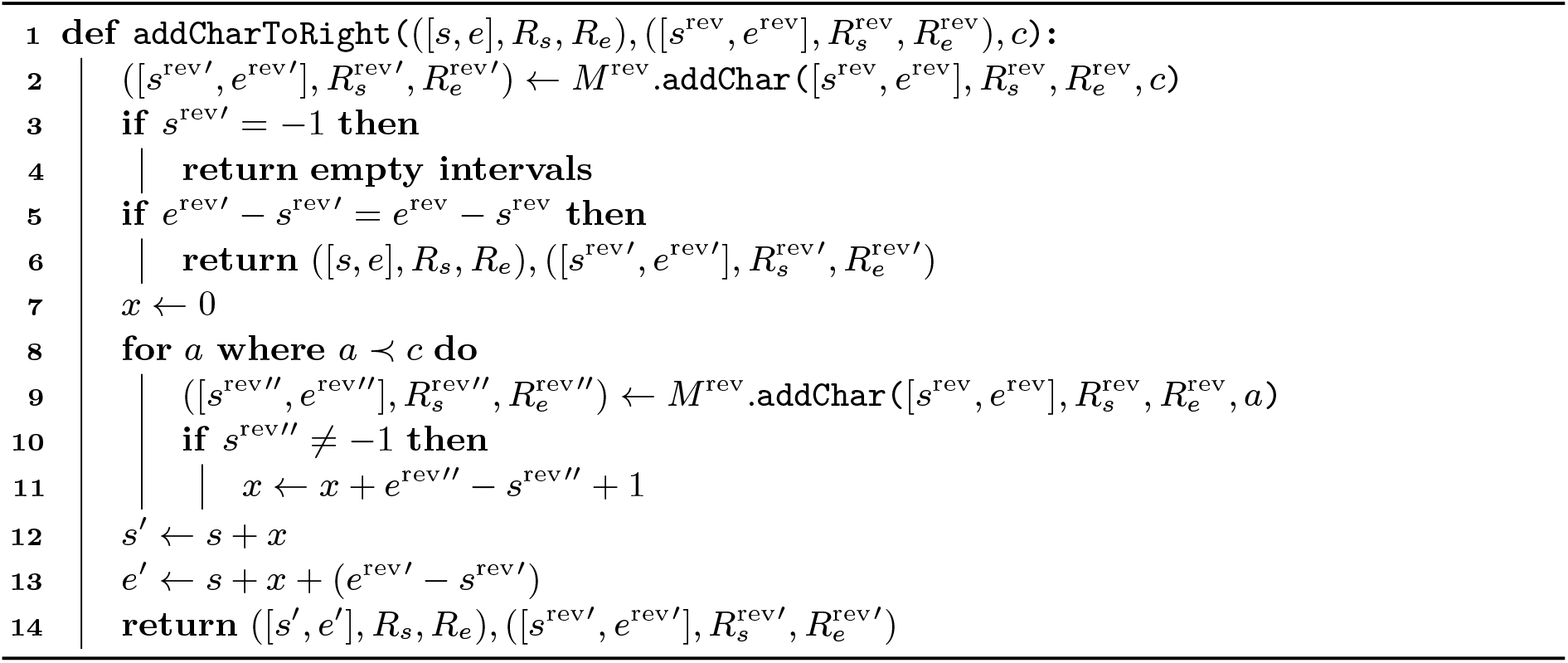

## Footnotes

1 Details available at https://github.com/biointec/b-move/tree/data/2024

2 Available at https://github.com/U-Ar/br-index

3 Available at https://github.com/biointec/columba

## Notes

https://github.com/biointec/b-move

### Competing Interest Statement

The authors have declared no competing interest.

## References

1 Omar Ahmed, Massimiliano Rossi, Sam Kovaka, Michael C. Schatz, Travis Gagie, Christina Boucher, and Ben Langmead. Pan-genomic matching statistics for targeted nanopore sequencing. iScience, 24(6):102696, 2021. doi:10.1016/j.isci.2021.102696.

2 Omar Y. Ahmed, Massimiliano Rossi, Travis Gagie, Christina Boucher, and Ben Langmead. Spumoni 2: improved classification using a pangenome index of minimizer digests. Genome Biology, 24(1):122, May 2023. doi:10.1186/s13059-023-02958-1.

3 Yuma Arakawa, Gonzalo Navarro, and Kunihiko Sadakane. Bi-Directional r-Indexes. In 33rd Annual Symposium on Combinatorial Pattern Matching (CPM 2022), volume 223 of Leibniz International Proceedings in Informatics (LIPIcs), pages 11:1–11:14, Dagstuhl, Germany, 2022. Schloss Dagstuhl – Leibniz-Zentrum für Informatik. doi:10.4230/LIPIcs.CPM.2022.11.

4 Andrej Baláž, Travis Gagie, Adrián Goga, Simon Heumos, Gonzalo Navarro, Alessia Petescia, and Jouni Sirén. Wheeler maps. In LATIN 2024: Theoretical Informatics, pages 178–192, Cham, 2024. Springer Nature Switzerland. doi:10.1007/978-3-031-55598-5_12.

5 Christina Boucher, Travis Gagie, I Tomohiro, Dominik Köppl, Ben Langmead, Giovanni Manzini, Gonzalo Navarro, Alejandro Pacheco, and Massimiliano Rossi. Phoni: Streamed matching statistics with multi-genome references. In 2021 Data Compression Conference (DCC), pages 193–202, 2021. doi:10.1109/DCC50243.2021.00027.

6 Nathaniel K. Brown, Travis Gagie, and Massimiliano Rossi. RLBWT tricks. In Data Compression Conference, DCC 2022, Snowbird, UT, USA, March 22-25, 2022, page 444. IEEE, 2022. doi:10.1109/DCC52660.2022.00055.

7 Michael Burrows and David Wheeler. A Block-Sorting Lossless Data Compression Algorithm. Research Report 124, Digital Equipment Corporation Systems Research Center, 130 Lytton Avenue, Palo Alto, California 94301, May 1994.

8 Dustin Cobas, Travis Gagie, and Gonzalo Navarro. A Fast and Small Subsampled R-Index. In 32nd Annual Symposium on Combinatorial Pattern Matching (CPM 2021), volume 191 of Leibniz International Proceedings in Informatics (LIPIcs), pages 13:1–13:16, Dagstuhl, Germany, 2021. Schloss Dagstuhl – Leibniz-Zentrum für Informatik. doi:10.4230/LIPIcs.CPM.2021.13.

9 The 1000 Genomes Project Consortium. A global reference for human genetic variation. Nature, 526(7571):68–74, Oct 2015. doi:10.1038/nature15393.

10 The Computational Pan-Genomics Consortium. Computational pan-genomics: status, promises and challenges. Briefings in Bioinformatics, 19(1):118–135, 10 2016. doi:10.1093/bib/bbw089.

11 Lore Depuydt, Luca Renders, Thomas Abeel, and Jan Fostier. Pan-genome de bruijn graph using the bidirectional fm-index. BMC Bioinform., 24(1):400, 2023. doi:10.1186/S12859-023-05531-6.

12 P. Ferragina and G. Manzini. Opportunistic data structures with applications. In Proceedings 41st Annual Symposium on Foundations of Computer Science, pages 390–398, 2000. doi: 10.1109/SFCS.2000.892127.

13 Travis Gagie, Gonzalo Navarro, and Nicola Prezza. Optimal-time text indexing in bwtruns bounded space. In Proceedings of the Twenty-Ninth Annual ACM-SIAM Symposium on Discrete Algorithms, SODA 2018, New Orleans, LA, USA, January 7-10, 2018, pages 1459–1477. SIAM, 2018. doi:10.1137/1.9781611975031.96.

14 Travis Gagie, Gonzalo Navarro, and Nicola Prezza. Fully Functional Suffix Trees and Optimal Text Searching in BWT-Runs Bounded Space. J. ACM, 67(1), jan 2020. doi:10.1145/3375890.

15 Adrián Goga, Lore Depuydt, Nathaniel K. Brown, Jan Fostier, Travis Gagie, and Gonzalo Navarro. Faster Maximal Exact Matches with Lazy LCP Evaluation. pages 123–132, 2024. doi:10.1109/DCC58796.2024.00020.

16 Gregory Kucherov, Kamil Salikhov, and Dekel Tsur. Approximate String Matching Using a Bidirectional Index. In Combinatorial Pattern Matching, pages 222–231, Cham, 2014. Springer International Publishing. doi:10.1007/978-3-319-07566-2_23.

17 T. W. Lam, Ruiqiang Li, Alan Tam, Simon Wong, Edward Wu, and S. M. Yiu. High Throughput Short Read Alignment via Bi-directional BWT. In 2009 IEEE International Conference on Bioinformatics and Biomedicine, pages 31–36, 2009. doi:10.1109/BIBM.2009.42.

18 Ben Langmead and Steven L. Salzberg. Fast gapped-read alignment with Bowtie 2. Nature Methods, 9(4):357–359, Apr 2012. doi:10.1038/nmeth.1923.

19 Heng Li and Richard Durbin. Fast and accurate short read alignment with Burrows–Wheeler transform. Bioinformatics, 25(14):1754–1760, 2009. doi:10.1093/bioinformatics/btp324.

20 Veli Mäkinen and Gonzalo Navarro. Succinct suffix arrays based on run-length encoding. In Combinatorial Pattern Matching, pages 45–56, Berlin, Heidelberg, 2005. Springer Berlin Heidelberg. doi:10.1007/11496656_5.

21 Udi Manber and Gene Myers. Suffix Arrays: A New Method for On-Line String Searches. SIAM Journal on Computing, 22(5):935–948, 1993. doi:10.1137/0222058.

22 Takaaki Nishimoto and Yasuo Tabei. Optimal-time queries on bwt-runs compressed indexes. In 48th International Colloquium on Automata, Languages, and Programming, ICALP 2021, July 12-16, 2021, Glasgow, Scotland (Virtual Conference), volume 198 of LIPIcs, pages 101:1–101:15. Schloss Dagstuhl - Leibniz-Zentrum für Informatik, 2021. doi:10.4230/LIPICS.ICALP.2021.101.

23 Luca Renders, Lore Depuydt, and Jan Fostier. Approximate Pattern Matching Using Search Schemes and In-Text Verification. In Bioinformatics and Biomedical Engineering, pages 419–435, Cham, 2022. Springer International Publishing. doi:10.1007/978-3-031-07802-6_36.

24 Luca Renders, Lore Depuydt, Sven Rahmann, and Jan Fostier. Automated design of efficient search schemes for lossless approximate pattern matching. In Research in Computational Molecular Biology, pages 164–184, Cham, 2024. Springer Nature Switzerland. doi:10.1007/978-1-0716-3989-4_11.

25 Luca Renders, Kathleen Marchal, and Jan Fostier. Dynamic partitioning of search patterns for approximate pattern matching using search schemes. iScience, 24(7):102687, 2021. doi: 10.1016/j.isci.2021.102687.

26 Massimiliano Rossi, Marco Oliva, Ben Langmead, Travis Gagie, and Christina Boucher. MONI: A pangenomic index for finding maximal exact matches, 2022. doi:10.1089/CMB.2021.0290.

27 Julian Seward. bzip2 and libbzip2 - a program and library for data compression. avaliable at http://www.bzip.org, 1996.

28 Mohsen Zakeri, Nathaniel K. Brown, Omar Y. Ahmed, Travis Gagie, and Ben Langmead. Movi: a fast and cache-efficient full-text pangenome index. bioRxiv, 2024. Accepted into RECOMB-seq 2024 proceedings track. doi:10.1101/2023.11.04.565615.

